# Genomic selection for any dairy breeding program via optimized investment in phenotyping and genotyping

**DOI:** 10.1101/2020.08.16.252841

**Authors:** Jana Obšteter, Janez Jenko, Gregor Gorjanc

## Abstract

This paper evaluates the potential of maximizing genetic gain in dairy cattle breeding by optimizing investment into phenotyping and genotyping. Conventional breeding focuses on phenotyping selection candidates or their close relatives to maximize selection accuracy for breeders and quality assurance for producers. Genomic selection decoupled phenotyping and selection and through this increased genetic gain per year compared to the conventional selection. Although genomic selection is established in well-resourced breeding programs, small populations and developing countries still struggle with the implementation. The main issues include the lack of training animals and lack of financial resources. To address this, we simulated a case-study of a small dairy population with a number of scenarios with equal resources yet varied use of resources for phenotyping and genotyping. The conventional progeny testing scenario had 11 phenotype records per lactation. In genomic scenarios, we reduced phenotyping to between 10 and 1 phenotype records per lactation and invested the saved resources into genotyping. We tested these scenarios at different relative prices of phenotyping to genotyping and with or without an initial training population for genomic selection. Reallocating a part of phenotyping resources for repeated milk records to genotyping increased genetic gain compared to the conventional scenario regardless of the amount and relative cost of phenotyping, and the availability of an initial training population. Genetic gain increased by increasing genotyping, despite reduced phenotyping. High-genotyping scenarios even saved resources. Genomic scenarios expectedly increased accuracy for young non-phenotyped male and female candidates, but also cows. This study shows that breeding programs should optimize investment into phenotyping and genotyping to maximise return on investment. Our results suggest that any dairy breeding program using conventional progeny testing with repeated milk records can implement genomic selection without increasing the level of investment.

## 1 Introduction

This paper evaluates the potential of maximizing genetic gain in dairy cattle breeding by optimizing investment into phenotyping and genotyping. Breeding programs strive to maximize genetic gain, which is a function of selection intensity, accuracy of selection, genetic variation, and generation interval. The conventional dairy breeding program uses an expensive and time-consuming progeny test. Genomic selection (Meuwissen et al., 2001; Schaeffer, 2006) achieves the same genetic progress faster and cheaper through a substantially reduced generation interval, increased accuracy of selection for young animals, and increased selection intensity of males (Schaeffer, 2006; Obšteter et al., 2019). Despite lower accuracy of sire selection compared to the conventional selection, genomic selection doubles the rate of genetic gain per year in dairy cattle (Wiggans et al., 2017).

All breeding programs operate with a limited resources allocated to breeding activities with the aim to maximize return on investment. Genomic selection is now a de-facto standard in leading breeding programs but is still challenging to implement in small national breeding programs or in developing countries. While leading breeding programs can service some small national breeding, developing countries require tailored breeding goals that will respond to the rapidly rising demand for a sustainable dairy production in local environment (Ducrocq et al., 2018; Marshall et al., 2019; Mrode et al., 2019). We hypothesize that these breeding programs need to evaluate priorities and could optimize the allocation of resources for phenotyping and genotyping to maximize return on investment. We base this hypothesis on the following simple examples (Table S1).

The accuracy of conventional (pedigree-based) estimates of breeding values increases with increasing heritability and increasing number of phenotype records per animal or its closest relatives (e.g. Mrode, 2005). For example, for a female-expressed trait with 0.25 heritability, the accuracy as a function of the number of repeated records per lactation (n) is 0.89 (n=10), 0.81 (n=5), 0.70 (n=2) and 0.62 (n=1). The corresponding accuracies for 100 sires tested on the total of 10,000 daughters are respectively 0.98 (n=10), 0.97 (n=5), 0.96 (n=2) and 0.93 (n=1).

The accuracy of genome-based estimates of breeding values similarly increases with increasing heritability and increasing number of phenotype records per genotyped animal. It also increases with increasing training population, decreasing genetic distance between training and prediction individuals, and decreasing number of effective genome segments (Daetwyler et al., 2008; Clark et al., 2011; Goddard et al., 2011). Following the previous example, assume 10,000 effective genome segments, 0.25 heritability, and a training population of 10,000 cows. The accuracy of genomic prediction for non-phenotyped animals as a function of the number of repeated records per lactation in the training population (n) is 0.76 (n=10), 0.71 (n=5), 0.63 (n=2), or 0.56 (n=1). These examples show diminishing returns with repeated phenotyping and a scope for optimizing return on investment in genomic breeding programs.

The accuracy of genomic prediction as a function of the number of genotyped and phenotyped cows (N) and the number of repeated records per lactation (n) is then 0.84 (N=20,000, n=5), 0.90 (N=50,000, n=2), or 0.93 (N=100,000, n=1). While these genomic prediction accuracies are lower than with progeny testing, shorter generation interval enables larger genetic gain per unit of time (Schaeffer, 2006).

Previous studies also explored the value of adding females to the training population (Van Grevenhof et al., 2012; Gonzalez-Recio et al., 2014). They concluded that accuracy has diminishing returns with increasing the number of genotyped and phenotyped animals in the training population, hence additional females are most valuable when training population is small. However, real breeding programs involve overlapping generations, individuals with a mix of phenotype, pedigree, and genotype information, various selection intensities, and other dynamic components. Thus, evaluating the optimal allocation of resources into phenotyping and genotyping is beyond these simple examples.

The above examples suggest that repeated phenotyping could serve as an internal financial reserve to enable dairy breeding programs to implement genomic selection and maximize return on investment. In dairy breeding the most repeatedly and extensively recorded phenotypes are milk production traits. There are different milk recording methods that differ in the recording frequency (International Committee for Animal Recording, 2017). The recording interval ranges from daily recording to recording every nine weeks, which translates to between 310 and 5 records per lactation. The different recording methods have different costs, which vary considerably between recording systems, countries, and even their regions. For example, some organizations require payment of a participation fee plus the cost per sample, while others include the fee in the sample cost, or cover the costs in other ways. There is also a huge variance in the way dairy breeding programs are funded. Some are funded by farmers and breeding companies while in some countries data recording or breeding are subsidized by state or charities. There are also situations where farmers do not record phenotypes and all genetic progress is generated in “research” nucleus herds.

The aim of this study was to evaluate the potential of maximizing genetic gain by optimizing investment into phenotyping and genotyping in dairy breeding programs. Since milk production traits are example of repeated phenotypes with diminishing returns, we aimed to optimize investment into milk recording and genotyping. To this end we have compared a dairy breeding program with conventional progeny testing and several genomic breeding programs under equal financial resources. To implement genomic selection, we reduced the number of milk records per lactation and invested the saved resources into genotyping. We compared the breeding programs in a case-study of a small cattle population where implementing genomic selection is challenging. The results show that reallocating a part of phenotyping resources to genotyping increases genetic gain regardless of the cost and amount of genotyping, and the availability of an initial training population.

## 2 Methods

The study aimed to evaluate the effect of different investment into phenotyping and genotyping with a simulation of a case-study of a small dairy breeding program. The simulation mimicked a real dairy cattle population of ~30,000 animals analyzed in Obšteter et al. (2019). Here we evaluated 36 genomic scenarios against the conventional scenario, all with equal available resources, but varying extent of phenotyping and genotyping. The conventional scenario implemented progeny testing and collected 11 phenotype records per lactation, while genomic scenarios reduced phenotyping and invested the saved resources into genotyping. The genomic scenarios differed in i) the number of phenotype records per lactation; ii) the relative cost of phenotyping and genotyping; and iii) the availability of an initial training population. All scenarios were compared on genetic gain and accuracy of selection.

### 2.1 Simulation of the base population, phenotype and historical breeding

The simulation mimicked a small dairy cattle breeding program of ~30,000 animals with ~10,500 cows. The introduction of effective genomic selection in such populations is challenging due to costs of assembling a training population and limited number of training animals. We use this population as a case-study to optimize investment into phenotyping and genotyping. The dairy breeding program aimed to improve production traits, which we simulated as a single polygenic trait. We used a coalescent process to simulate genome comprised of 10 cattle-like chromosomes, each with 10^8^ base pairs, 1,000 randomly chosen causal loci, and 2,000 randomly chosen marker loci. We sampled the effects of causal loci from a normal distribution and use them to calculate animal’s breeding value (*a_i_*) for dairy performance (*y_ijkl_*), which was affected also by a permanent environment (*p_i_*) herd (*h_j_*), herd-year (*hy_jk_*), herd-test-day (*htd_jkl_*), and residual environment (*e_ijkl_*) effects: y_ijkl_ = a_i_ + p_i_ + h_j_ + hy_jk_ + htd_jkl_ + e_ijkl_.

We sampled permanent environment effects from a normal distribution with zero mean and variance equal to the base population additive genetic variance (*σ^2^_a_*). We sampled herd, herd-year, and herd-test-day effects each from a normal distribution with zero mean and variance of 1/3 *σ^2^_a_*. Finally, we sampled residual environment effects from a normal distribution with zero mean and variance of *σ^2^_a_*. This sampling scheme gave a trait with 0.25 heritability and 0.50 repeatability. With the simulated genome and phenotype architecture we have initiated a dairy cattle breeding program and ran it for 20 years of conventional selection with progeny-testing based on 11 cow phenotype records per lactation. The detailed parameters of the simulation are described in Obšteter et al. (2019). In summary, in the breeding program we selected 3,849 out of 4,320 new-born females. We selected 139 bull dams out of cows in the second, third, and fourth lactation. We generated 45 male calves from matings of bull dams and progeny tested sires (elite matings). Out of these we chose 8 for progeny testing of which 4 were eventually selected as sires for widespread insemination of cows. We made all selection decisions based on pedigree-based estimates of breeding values. The 20 years represented historical breeding and provided a starting point for evaluating future breeding scenarios, which we ran for additional 20 years.

### 2.2 Scenarios

We evaluated 36 genomic scenarios with varying extent of phenotyping and genotyping against the conventional scenario, all with equal resources. The conventional scenario continued the breeding scheme from the historical breeding. It used progeny testing and 11 phenotype recordings per lactation (named C11), corresponding to the standard ICAR recording interval of 4 weeks (International Committee for Animal Recording, 2017). We assumed that this scenario represented the total resources for generating data. We then created genomic scenarios by distributing resources between phenotyping and genotyping - we reduced phenotyping and invested the saved resources into genotyping. In genomic scenarios we selected females as in the conventional scenario and males on genomic estimates of breeding values. The number of genomically tested male candidates varied according to the genotyping resources in a specific scenario. We selected 5 males with the highest genomic estimates of breeding value as sires for widespread insemination of cows. We evaluated the genomic scenarios with a varying number of phenotype records per lactation, relative cost of phenotyping to genotyping, and the availability of an initial training population.

In genomic scenarios we reduced the number of phenotype records per lactation to between 10 and 1. Three of them followed ICAR standards of 9, 8, and 5 records per lactation, corresponding to recording intervals of 5, 6, and 9 weeks. Additionally, we tested the non-standard 10, 2, and 1 records per lactation. We named the scenarios as “GX” with X being the number of records per lactation. Genomic scenarios next varied the relative cost of phenotyping ($P) to genotyping ($G). We compared the cost of 11 phenotype records per lactation to the cost of one genotype. By increasing the number of records per lactation the cost per record decreased by 6%. The cost of genotyping remained constant. Based on a survey of several breeding programs, milk recording organizations, and genotyping providers we have considered three cost ratios of $P:$G: 2:1, 1:1, and 1:2. The reduction in phenotyping and the relative cost of phenotyping to genotyping dictated the number of genotyped animals (Table 1). For example, assume that the cost per milk record is €1.78 with 11 records per lactation (in total €19.55). Further assume that the cost per milk record is €1.89 with 10 records per lactation (in total €18.91). This second scenario saves €0.64 that can be used for genotyping. Applying this to our simulated populations with 10,852 active cows we could genotype 356 animals every year when $P:$G = 1:1.

**Table 1:** Number of genotyped animals per year by scenario and relative cost of phenotyping to genotyping ($P:$G). Scenarios are named “G” for genomic, followed by the number of phenotype records per lactation. For the $P:$G we compared the cost of 11 phenotype records per lactation to the cost of one genotype. The number of phenotype records and the $P:$G dictated the number of genotyped animals. We genotyped females (F) and males (M) in 7:1 ratio. We genotyped the females to update and increase the training population and males for selection.

| Relative cost | Scenario |  |  |  |  |  |
| --- | --- | --- | --- | --- | --- | --- |
|  | G10 | G9 | G8 | G5 | G2 | G1 |
| \$P:\$G = 1:2 | 160 F | 350 F | 590 F | 1,610 F | 3,230 F | 3,850 F |
|  | 22 M | 50 M | 85 M | 235 M | 465 M | 565 M |
| \$P:\$G = 1:1 | 310 F | 700 F | 1,180 F | 3,230 F | 3,850 F | 3,850 F |
|  | 45 M | 100 M | 165 M | 465 M | 925 M | 1,125 M |
| \$P:\$G = 2:1 | 620 F | 1,400 F | 2,360 F | 3,850 F | 3,850 F | 3,850 F |
|  | 90 M | 200 M | 335 M | 925 M | 1,845 M | 2,245 M |

We invested the saved resources into genotyping females and males in ratio 7:1 based on our previous work (Obšteter et al., 2019). We genotyped first parity cows. This maximized the accuracy of genomic prediction, since it reduced genetic distance between training and prediction population, prevented the loss of investment with culled heifers, and minimized the time to obtain a phenotype linked to a genotype. To maximize the genetic gain, we genotyped male calves from elite matings and other high parent average matings. In scenarios where the resources for genotyping females were larger than the cost of genotyping all first parity cows, we did not reallocate the excess of resources to genotyping males for consistency (we saved resources in those cases).

Lastly, we created scenarios with and without an initial training population for genomic prediction. When we assumed an initial training population was available, we genotyped all active cows (10,852) and progeny tested sires (100) before the first genomic selection of males. When an initial training population was not available, we yearly genotyped a designated number of first parity cows until the training population reached 2,000 cows. Once we reached this goal, we started to genotype both females and males as specified in Table 1. At that point we started genomic selection of males.

### 2.3 Estimation of breeding values

We selected animals based on their breeding values estimated from a pedigree or single-step genomic repeatability model with breeding value, permanent environment, and herd-year as random effects. We did not fit the herd-test-day effect as data structure of this small population did not enable its accurate estimation. We estimated breeding values once a year with blupf90 (Misztal et al., 2018) with default settings. In the estimation we included all available phenotype and pedigree records for all active, phenotyped, or genotyped animals, and additional three generations of their ancestors. We used at most 25,000 genotype records due to a limit in the academic software version. When we accumulated more than 25,000 genotyped animals, we removed genotypes of the oldest animals in favor of the latest genotyped cows and male selection candidates.

### 2.4 Analysis of scenarios

All scenarios had equal resources. We compared the scenarios based on their final genetic gain, which indicated return on investment, accuracy of selection, and selection intensity. We measured the genetic gain as an average true breeding value by year of birth and standardized it to have zero mean and unit standard genetic deviation in the first year of comparison. We measured the accuracy of breeding values as the correlation between true and estimated breeding values. We measured the accuracy separately for four groups of animals: i) male candidates (genotyped and non-phenotyped); ii) sires (currently used in artificial insemination); iii) female candidates (non-genotyped and non-phenotyped); and iv) cows (all active phenotyped cows and bull dams). We computed the intensities of two-stage selection (*i*) by integrating standard bivariate distribution 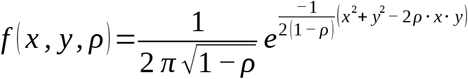 as 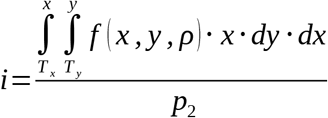, where *x* is the is the parent average, *y* is the genomic breeding value, and *ρ* is the correlation between the variables (Young, 1964; Jopson et al., 2004). We computed the correlation *ρ* by dividing the scenario-specific accuracy of parent average by the accuracy of genomic prediction. T_x_ is the standardized cut-off for the proportion of all new-born male calves selected for genomic testing (*p_i_*) based on parent average in the first selection stage. T_y_ is the standardized cut-off for the proportion of all newborn males selected as sires (*p_2_*) based on genomic breeding values in the second selection stage. We repeated simulation of the base population and each scenario 10 times and summarized them with mean and standard deviation across the replicates. We used Tukey’s multiple comparison test to test the significance of the difference between means.

## 3 Results

Genomic scenarios increased the genetic gain compared to the conventional scenario regardless of the number of phenotype records per lactation, relative cost of phenotyping to genotyping, and the availability of an initial training population. Genomic scenarios with an initial training population increased the genetic gain of the conventional scenario by up to 143%, despite reduced phenotyping. Genetic gain increased with increasing investment into genotyping. Genomic scenarios increased accuracy for non-phenotyped male and female candidates, and cows. Scenarios without an initial training population showed the same trends for genetic gain and accuracy. Although these scenarios had a slightly smaller genetic gain due to delayed implementation of genomic selection, they still increased the genetic gain of the conventional scenario by up to 134%. We present these results in more details in the following sub-sections separately for settings with and without an initial training population available.

### 3.1 Genetic gain with an initial training population

Genomic scenarios with an initial training population increased the genetic gain of the conventional scenario using the same resources. The genetic gain increased with increasing investment in genotyping, despite reduced phenotyping (Figure 1 and Table S2). We show the corresponding intensities of sire selection in Table S3. In the $P:$G = 1:1 setting, the genomic scenarios increased the genetic gain of the conventional scenario between 79% and 143%. By reducing the number of phenotype records from 11 (C11) to 10 per lactation (G10), we saved resources for genotyping 355 animals per year (45 male candidates). This small change increased male selection intensity from 0.10 to 0.26 and coupled with a shorter generation interval increased the genetic gain by 79% (from 3.01 to 5.41). By reducing the number of phenotype records to nine or eight per lactation (G9 or G8), we respectively saved resources to genotype 800 or 1,345 animals per year (100 or 165 male candidates). This respectively increased male selection intensity to 0.39 or 0.49, and genetic gain by 109% or 120% (from 3.01 to 6.30 or 6.62). We achieved the highest genetic gain, between 135% and 143% of the conventional scenario (between 7.07 and 7.33), when we collected between five and one phenotype records per lactation. In these three scenarios we saved resources for genotyping between 3,230 and 3,850 (all) cows and between 465 and 1,125 male candidates per year, and achieved male selection intensity between 0.77 and 0.92.

**Figure 1.**
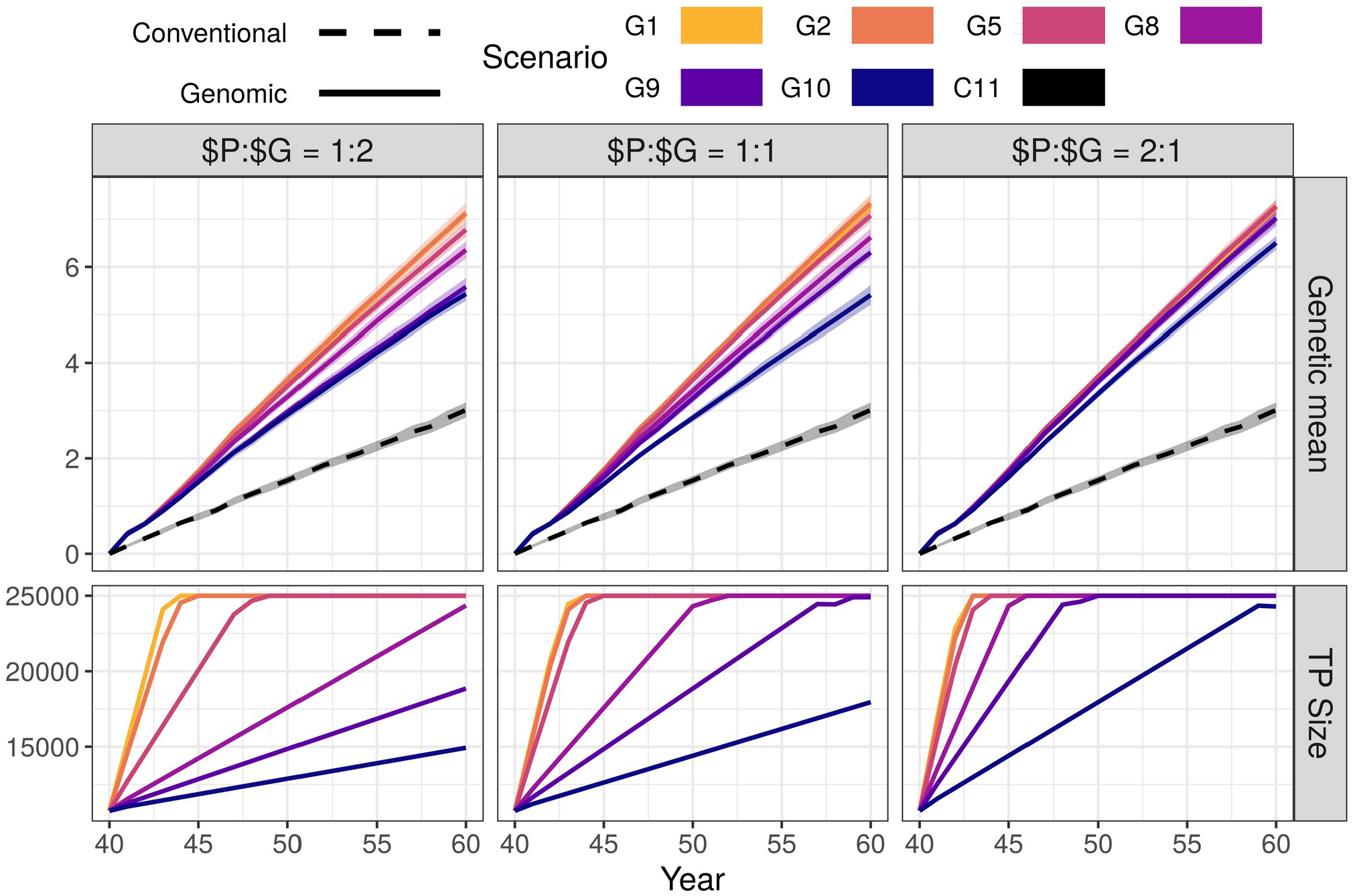
Genetic gain and training population size by scenario and relative cost of phenotyping to genotyping ($P:$G) with an initial training population (TP). The figure presents the means (lines) and 95% confidence intervals (polygons) across 10 replicates for the conventional (C) and genomic (G) scenarios, with numbers indicating the number of phenotype records per lactation. For the $P:$G we compared the cost of 11 phenotype records per lactation to the cost of one genotype.

Changing the relative cost of phenotyping to genotyping did not change the trend in genetic gain. In the G10 scenario of the $P:$G = 1:2 setting we yearly genotyped 182 animals (22 male candidates) and increased the genetic gain by 80% (from 3.01 to 5.43). In the G10 scenario of the $P:$G = 2:1 setting we yearly genotyped 710 animals (90 males candidates) and increased the genetic gain by 116% (from 3.01 to 6.50). When we maximized the investment into genotyping (G1), we genotyped between 565 and 2,245 male candidates and all females. This achieved a comparable genetic gain, between 136% and 143% of the conventional scenario, regardless of the relative cost of phenotyping to genotyping and male selection intensities.

The high-genotyping scenarios achieved the observed genetic gain without using all the resources (marked bold in Table S2). In these scenarios the resources designated to genotyping females exceeded the cost of genotyping all females. These savings could cover additional 11 phenotype records per lactation for between 169 and 5,950 animals, or between 85 and 11,900 additional genotypes.

In Figure 1 we also show the growth of the training population for genomic prediction. The training population started with ~10,000 individuals and grew until reaching 25,000 individuals. The increase was not linear through all generations, since the procedure for choosing the training animals changed when the training population exceed 25,000 (only latest females and male candidates included)

### 3.2 Accuracy with an initial training population

Compared to the conventional scenario, genomic scenarios increased accuracy for young non-phenotyped and genotyped male, non-phenotyped and non-genotyped female candidates, and cows, but decreased accuracy for sires (Figure 2 and Table S4). In the $P:$G=1:1 setting, the accuracy for young genomically tested male candidates ranged between 0.90 and 0.91 regardless of the amount of phenotyping and genotyping. This accuracy was between 0.53 and 0.54 higher compared to the pre-selection for progeny testing and between 0.03 and 0.04 lower compared to the sire selection in the conventional scenario. In contrast, the accuracy for already selected sires decreased with decreasing investment into phenotyping and was between 0.11 and 0.23 lower than in conventional scenario. We observed the lowest accuracy for sires (0.63) when we invested the most into genotyping (G1) and the highest (0.75) when we invested the most into phenotyping (G10).

**Figure 2.**
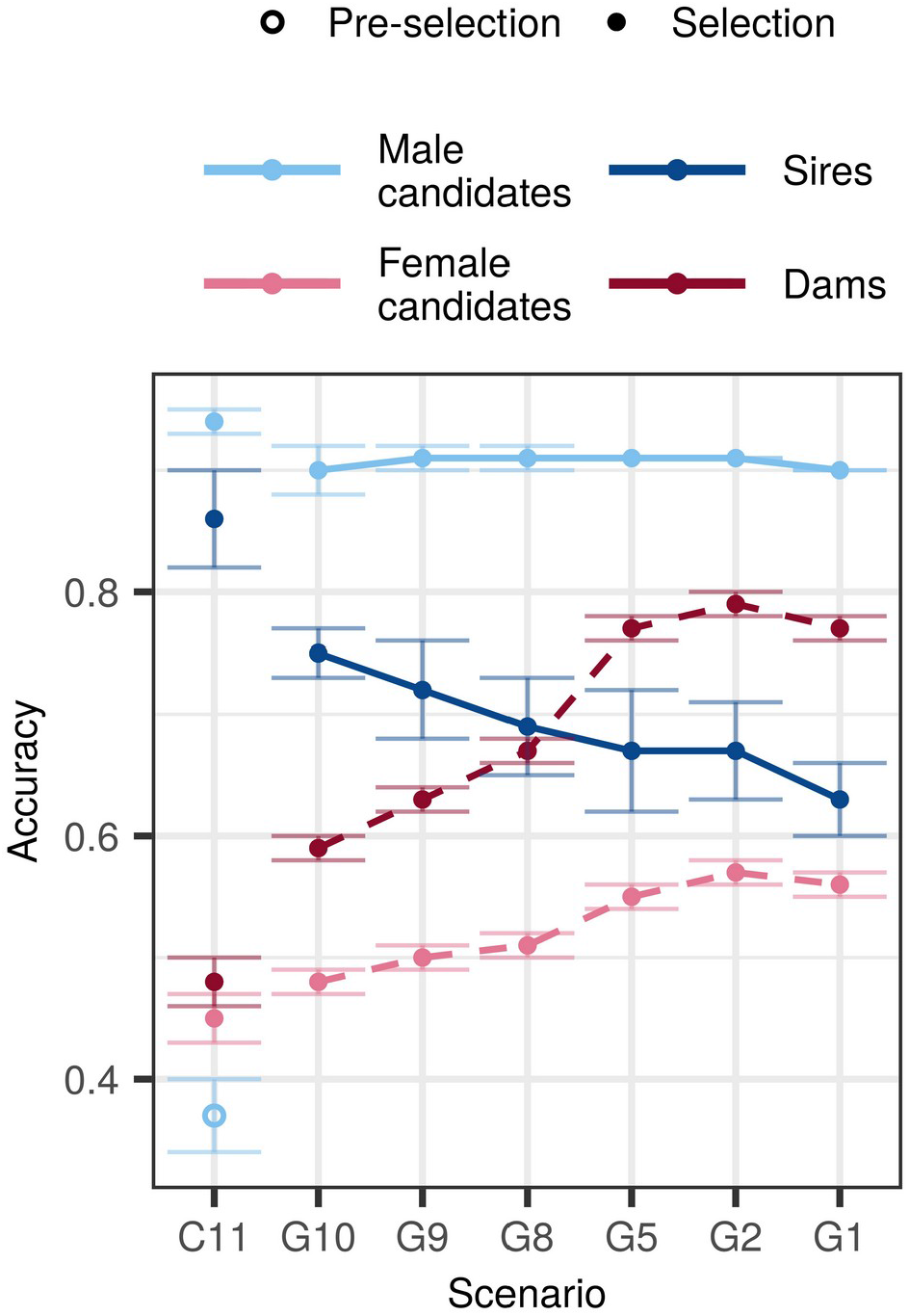
Accuracy by scenario with an initial training population and equal cost of phenotyping and genotyping. The figure presents the means (lines) and 95% confidence intervals (error bars) across 10 replicates for the conventional (C) and genomic (G) scenarios with numbers indicating the number of phenotype records per lactation. The cost of phenotyping was defined as the cost of 11 phenotype records per lactation, and the cost of genotyping as the cost of one genotype. Conventional selection implemented two-stage selection for males, hence we present the accuracy of pre-selection for progeny testing (empty point) and the accuracy of sire selection (solid point).

The accuracy for non-genotyped female candidates and cows increased with increasing genotyping, despite reduced phenotyping. We observed the highest accuracy for female candidates (0.55 - 0.57) and cows (0.77 - 0.79) when we recorded between five and one phenotype record per lactation and invested the rest into genotyping. Compared to the conventional scenario, the genomic scenarios increased the accuracy between 0.03 and 0.11 for female candidates, and between 0.11 and 0.29 for cows.

Changing the relative cost of phenotyping to genotyping affected primarily the accuracy for female candidates and cows. In the majority of scenarios the accuracy increased with decreasing the relative cost of genotyping, which enabled more genotyping. We observed the largest difference of 0.06 for female candidates and 0.12 for cows when we changed the relative cost of phenotyping from half to twice the cost of genotyping. Changing the relative costs, however, did not change the accuracy trends.

### 3.3 Scenarios without an initial training population

#### 3.3.1 Genetic gain

Genomic scenarios without an initial training population also increased the genetic gain of the conventional scenario using the same resources. The trends were in line with what we observed with an initial training population, that is, increasing genotyping increased genetic gain despite reduced phenotyping (Figure 3). However, all corresponding scenarios achieved between 2% and 28% smaller genetic gain than when an initial training population was available (Table S2).

**Figure 3.**
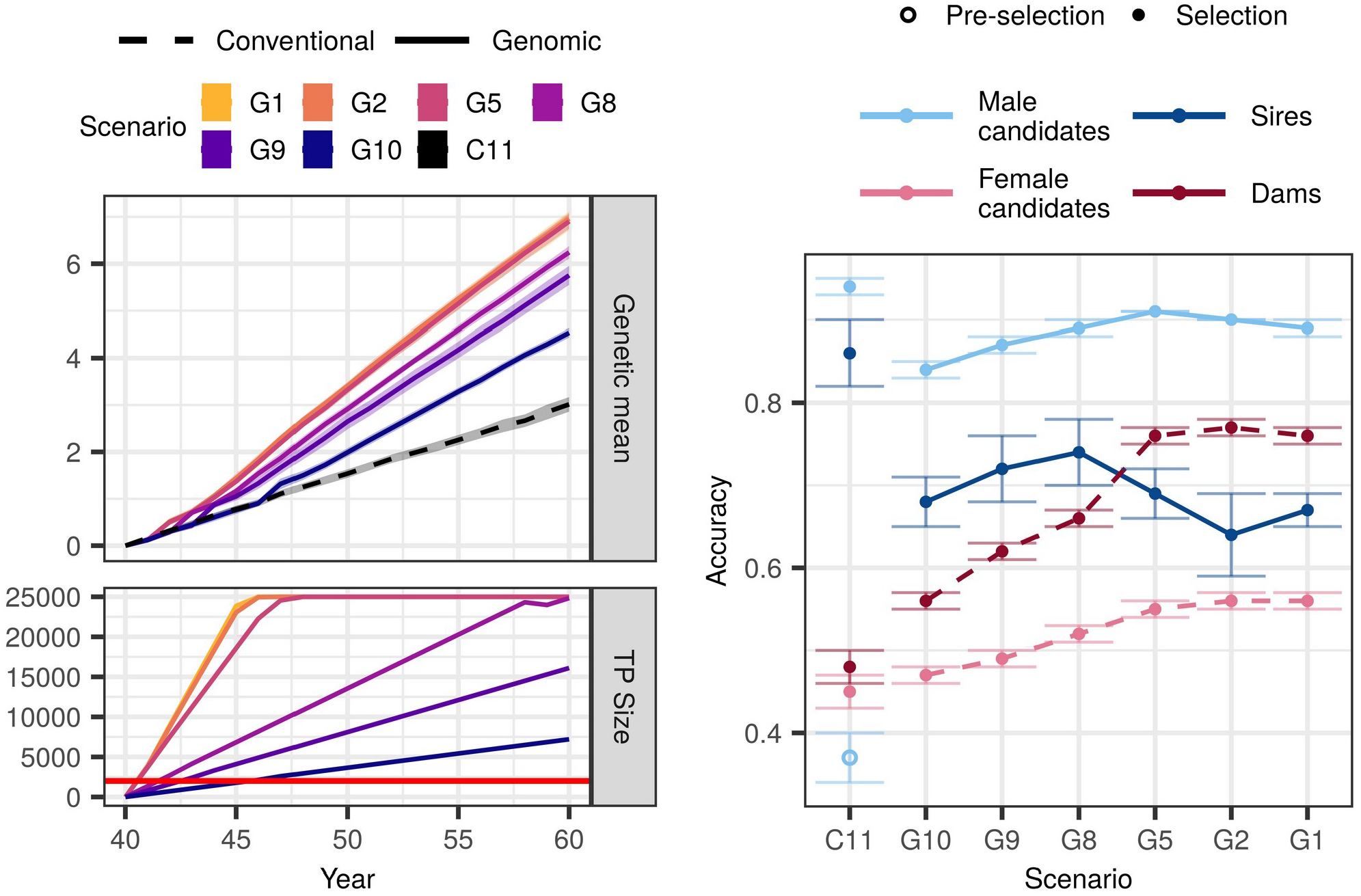
Genetic gain, training population size, and accuracy by scenario without an initial training population (TP) and equal cost of phenotyping and genotyping. The figure presents the means (lines or points) and 95% confidence intervals (polygons or error-bars) across 10 replicates for the conventional (C) and genomic (G) scenarios with numbers indicating the number of phenotype records per lactation. The cost of phenotyping was defined as the cost of 11 phenotype records per lactation, and the cost of genotyping as the cost of one genotype.The red line marks the condition of required 2,000 training animals to start genomic selection. Conventional selection implemented two-stage selection for males, hence we present the accuracy of the pre-selection for progeny testing (empty point) and the accuracy of sire selection (solid point).

In the $P:$G setting, genomic scenarios increased the genetic gain of the conventional scenario between 51% and 131%. Compared to when we had an initial training population, the corresponding scenarios achieved between 2% and 16% lower genetic gain. This difference was the largest when we invested the least into genotyping (G10). In this scenario we needed six years to build a training population of 2,000 cows and implement genomic selection, since we only genotyped 355 cows per year. We observed the smallest difference in the scenario that collected two phenotype records per lactations (G2) and implemented genomic selection in the first year.

Changing the relative cost of phenotyping to genotyping did not change the overall trend. In the $P: $G=1:2 setting, the genomic scenarios increased genetic gain of the conventional scenario between 31% and 126%. That was between 4% and 28% less than corresponding scenarios with an initial training population. In the $P:$G=2:1 setting, the genomic scenarios increased the genetic gain of the conventional scenario between 86% and 134%, which was between 3% and 14% less than corresponding scenarios with an initial training population.

#### 3.3.2 Accuracy

Similar to the scenarios with an initial training population, genomic scenarios without it increased the accuracy for non-phenotyped male and female candidates, and cows (Figure 3 and Table S4). In the $P:$G=1:1 setting the accuracy for male candidates ranged between 0.84 and 0.91. In contrast to scenarios with an initial training population, the accuracy increased with increasing investment into genotyping. The accuracy for sires ranged between 0.64 and 0.74. Contrary to when we had an initial training population, we observed no clear trend of either increasing or decreasing accuracy with decreasing investment into phenotyping. For female candidates the accuracy ranged between 0.47 and 0.56, and for cows between 0.56 and 0.76. For both the accuracies followed the trends of when we had an initial training population - increasing genotyping increased the accuracy.

Changing the relative cost of phenotyping to genotyping affected the accuracy for non-genotyped female candidates, cows and male candidates. Decreasing the relative cost of genotyping to phenotyping increased the accuracy in the majority of the scenarios, particularly the low-genotyping ones.

## 4 Discussion

Our results show that any dairy breeding program using conventional progeny testing with repeated milk records can implement genomic selection without extra costs. While breeding programs have established funding for phenotyping, not all of them have established funding for genotyping. We show that by reallocating a part of phenotyping resources into genotyping, breeding programs can implement genomic selection and substantially increase genetic gain regardless of the amount and cost of genotyping, and availability of an initial training population. However, increasing investment in genotyping has diminishing returns, which suggests that breeding programs should optimize the investment into phenotyping and genotyping to maximize return on investment for selection and management. The results raise four discussion points: 1) how optimizing the investment in phenotyping and genotyping affects genetic gain; 2) how optimizing the investment in phenotyping and genotyping affects accuracy; 3) implications for dairy breeding programs; and 4) limitations of the study. We first discuss the results under equal cost of phenotyping and genotyping, and an initial training population available. We then discuss changes at different costs and no initial training population.

### 4.1 Genetic gain with an initial training population

#### 4.1.1 Genomic vs. conventional selection

Implementing genomic selection by optimizing the investment in phenotyping and genotyping increased genetic gain compared to the conventional selection, mainly due to reduced generation interval in sire selection paths. This improvement is in agreement with previous theoretical studies (Schaeffer, 2006; Pryce et al., 2010; Obšteter et al., 2019). Empirical studies confirm this; in the US Holstein population the generation interval for the sires of sires and sires of dams paths recently decreased between 25% and 50% compared to the conventional selection (García-Ruiz et al., 2016). Van Grevenhof et al. (2012) also showed that when genomic selection halves the generation interval, a training population with ~2,000 individuals with own performance or ~3,500 individuals with ten progeny gives comparable response as conventional selection for a trait with intermediate heritability.

Another major advantage of the genomic scenarios was increased intensity of sire selection. A costly and lengthy progeny-testing limits the number of tested male candidates in conventional selection. Genomic selection significantly reduces the cost of testing (Schaeffer, 2006) and thus allows for testing more male candidates. In the US Holstein population, genomic selection improved the selection differential for all traits, particularly for traits with low heritability, such as health and fertility (García-Ruiz et al., 2016).

#### 4.1.2 Increasing the investment into genotyping

Genetic gain increased with increased investment into genotyping. This was mainly due to higher intensity of sire selection, since more resources for genotyping allowed us to test more male candidates while selecting the same number. A larger investment into genotyping also increased the update and total size of the training population, which increased the accuracy of female selection (we discuss this in the next sub-section).

The genetic gain had diminishing relationship with investment into genotyping. This has important implications for dairy breeding programs, since they use phenotypes also for management, and we discuss this separately. The results showed that investing resources of more than six phenotype records into genotyping did not significantly improve the genetic gain. There are three reasons for this. First, increasing female training population has diminishing relationship with genetic gain (Van Grevenhof et al., 2012; Gonzalez-Recio et al., 2014). Since scenarios with an initial training population started with ~10,000 genotyped and phenotyped cows, enlarging the training population had a marginal effect. Consequently, the accuracy of sire selection in genomic scenario was high regardless of the amount of genotyping. Second, increasing investment into genotyping did not proportionally increase the size of the training population due to the limit of 25,000 animals of the training population. And third, the intensity of sire selection had diminishing return with increasing number of genotyped male candidates. This agrees with Reiner-Benaim et al. (Reiner-Benaim et al., 2017) that showed an increased genetic gain with increasing the number of tested male candidates, but with a diminishing return. While they achieved the maximum profit with four selected sires out of 1,721 tested candidates, they achieved 99% or 90% of the maximum profit with respectively 740 or 119 tested candidates. The same three reasons enabled comparable maximum genetic gain regardless of the relative price of phenotyping to genotyping. In general, selecting less than 2% of the tested males and updating the training population with at least 35% of first-parity cows resulted in the maximum genetic gain.

While genetic gain increases with the number of cows in training population, it does not increase with the number of repeated records. The scenarios with the largest genetic gain therefore had a training population with many cows and few repeated records (Figure S1). However, since we used the single-step genomic prediction, the phenotypes of the non-genotyped animals contributed to the estimation as well. Effectively, all scenarios thus operated with the same number of phenotyped animals.

We should emphasize, that some of the high-genotyping scenarios achieved the observed genetic gain at a lower total cost, since they could not use all the saved resources for genotyping females in the studied population. The saved resources could be invested back into phenotyping females for milk production or novel traits, genotyping more male candidates, or other breeding actions.

### 4.2 Accuracy with an initial training population

Despite reduced phenotyping, genomic scenarios increased the accuracy for young non-phenotyped calves and cows. When the accuracy of parent average is already high, genomic prediction increases primarily the accuracy of the Mendelian sampling term. But when the accuracy of parent average is low, such as for the offspring of parents with little or no own or progeny information, genomic information increases accuracy both for the parent average and the Mendelian sampling term (Daetwyler et al., 2007; Wolc et al., 2011).

#### 4.2.1 Accuracy for males

For male candidates, genomic prediction more than doubled the accuracy compared to the parent average used for pre-selection of male calves for progeny testing. This is in agreement with two-fold accuracy increase in dairy (Schaeffer, 2006) and layers (Wolc et al., 2011). Within the genomic scenarios, the accuracy for male candidates was high regardless of the amount of genotyping and phenotyping for two reasons. First, the accuracy of their parent average was high, since we tested offspring of elite matings. Second, starting with an initial 10,000 training population gave an adequate accuracy that was additionally boosted by using all available information jointly through the single-step genomic prediction. Using single-step genomic prediction also removed the bias due to pre-selection (Jibrila et al., 2020).

In contrast, reducing phenotyping decreased the accuracy of already selected sires. We believe this is due to two reasons. First, since sires are the very best animals, their breeding values are close together in the tail of the distribution. Due to small differences between the sires, each additional phenotypic record helps to differentiate them and thus increases the accuracy. Second, as we invested more into genotyping, the training population grew quicker and reached the limit of 25,000. At this point we removed sires’ in favor of cows’ genotypes, hence prediction for sires depended only on daughters’ data and no longer on their own genotype. However, since this is the accuracy after the selection has already been made, it is not of great interest for breeding.

#### 4.2.2 Accuracy for females

Genomic scenarios increased the accuracy for cows compared to the conventional scenario. Besides increasing the accuracy of Mendelian sampling term, using genomic information increases genetic connectedness between individuals from different management units (Yu et al., 2017; Powell et al., 2019). Increased connectedness in turn increases the accuracy of prediction regardless of the heritability and the number of causal loci or markers (Yu et al., 2018). This improvement is important because we selected bull dams for elite mating from cows.

The accuracy for cows increased with increasing investment into genotyping, despite reduced phenotyping due to three reasons. First, more cows had both genomic and phenotypic information, which increased the accuracy of their estimated breeding values. Second, more genotyped cows increased genetic connectedness (Yu et al., 2018). And third, investing more into genotyping translated into larger training population and its yearly update. As shown by previous studies (Van Grevenhof et al., 2012; Gonzalez-Recio et al., 2014), the accuracy of genomic prediction increases with increasing the size of a female training population. They showed that the accuracy of 0.70 is achieved with ~20,000 animals as in our study. Same studies shown that, as with genetic gain, accuracy had a diminishing return with the size of the training population. We observed plateau in accuracy when we invested more than six phenotype records (out of eleven) into genotyping.

Accuracy for female candidates followed the trend for cows, but at lower values. Female candidates were not genotyped nor phenotyped, hence their accuracy mainly reflected the accuracy of their parent average. Increasing genotyping increased the accuracy for cows and in turn increased the accuracy of female candidate’s parent average. The benefit of this increase was not large, since the intensity of female selection was low. However, there is potential for this benefit to be larger with sexed semen and embryo transfer.

### 4.3 Scenario without an initial training population

#### 4.3.1 Genetic gain

We also considered that some populations do not have access to an initial training population and have to initialize one themselves. These genomic scenarios still increased genetic gain compared to the conventional scenario, but achieved lower genetic gain than corresponding scenarios with an initial training population available. Increasing the investment into genotyping compensated for starting without a training population in two ways. First, it shortened the time to obtain the targeted 2,000 genotypes required to implement genomic selection down to one year in high-genotyping scenarios. Second, it shortened the time to build a training population in which an additional record had negligible effect on accuracy (Gonzalez-Recio et al., 2014).

#### 4.3.2 Accuracy

Accuracy in scenarios without an initial training population closely followed the trends of the corresponding scenarios with an initial training population available. We observed minor differences in the low genotyping scenarios that had reduced accuracy for male candidates and sires. We attribute this to a smaller training population. Buch et al. (2012) showed that for new traits and large-scale recording, we can achieve 75% of the maximum genomic accuracy within first two to three years of recording. In our study we shortened this period even more by including the historical data through the single-step genomic prediction.

### 4.4 Implications

The results suggest that any dairy breeding program using conventional progeny testing with repeated milk records can implement genomic selection without extra costs by optimizing the investment of resources into breeding actions. Here we propose funding the genotyping with a part of resources for milk recording, since we can manipulate the number of repeated records. Breeding programs could reduce phenotyping for a different trait that they record repeatedly and is perhaps less crucial for management. They could also reallocate the funds from another breeding action.

Additionally, we could optimize which individuals to genotype and phenotype, as well as the computational costs. Selective phenotyping was shown to increase the accuracy of genomic prediction up to 20% with small sample sizes in plant breeding (Heslot & Feoktistov, 2017; Akdemir & Isidro-Sánchez, 2019). Similarly, selective genotyping of cows from the distribution tails has been shown to increase the accuracy of genomic prediction by 15% (Jenko et al., 2017). We expect this would further increase the return on investment, but increase the complexity of optimization. Regarding computing costs, the problem of a large number of genotypes can be alternatively solved by using methods with reduced computational costs. Examples of such methods are the algorithm for proven and young (Misztal et al., 2014) and singular value decomposition of the genotype matrix (Ødegård et al., 2018). Also, as shown in our study, we can achieve large genetic gain with a relatively small training population of recent genotypes.

The economic efficiency of breeding programs strongly depends on which stakeholders fund which breeding action. Different programs have different investment schemes, often complex. The scenarios presented in this paper are of little value for programs where funding for phenotyping and genotyping is disconnected. Similarly, optimizing the investment into phenotyping is not of interest for breeding programs with abundant use of automated milking systems where the cost of phenotyping does not depend on the number of records. However, in populations with small herds the use of automated milking systems is limited. Further on, genomic selection could benefit some settings more than others. For example, genomic information is especially important for generating genetic connectedness in systems with small herd sizes, geographically dispersed farms, and limited use of artificial insemination, often found in low-to mid-income countries (Powell et al., 2019). The same benefits are expected for small ruminant programs that do not actively exchange sires between herds (Kasap et al., 2018).

High cost of genotyping diminishes the benefit of the proposed solutions. The relative cost of phenotyping to genotyping at which genomic selection is not beneficial to conventional selection depends on a number of factors: i) the number of females in the recorded population, since it dictates the savings from reducing the phenotyping; ii) the number of phenotype records the breeders are willing to sacrifice; iii) the availability of an initial training population; and iv) the ratio of genotyped males and females. In our case, if genotyping was ten-times more expensive than phenotyping with 11 records, we would save resources to genotype between 36 (ten phenotypic records) and 900 (one phenotypic record) animals. While such numbers of genotyped animals could allow for efficient genomic selection if we had an initial training population and genotyped only males, it would probably not be viable if we had to build the training population ourselves.

We did not account for the benefits of genotyping besides genomic selection. Genomic information has additional value for i) parentage verification, parentage discovery, or correction of parentage errors; ii) management of monogenic diseases and traits, which can prevent economic losses caused by spreading lethal alleles or create economic gain by adding value to the products; iii) better monitoring and control of inbreeding (Sonesson et al., 2012) and optimization of matings (Obšteter et al., 2019); iv) determination of animals’ breed composition, which serves to identify the most appropriate breed or cross-breed in a production system, or improve the structure of the training population (Marshall et al., 2019). Additional uses of genotypes increase the return on investment beyond what we measured in this study.

### 4.5 Limitations of the study

#### 4.5.1 Reducing the number of phenotype records

Balancing phenotyping and genotyping can lead to conflicts between managing production (shortterm goal) and achieving genetic gain (long-term goal). Producers use phenotype records to manage animals’ health, reproduction and feed composition, which affect the success of dairy cattle production in the short-term. Besides managing production, milk recording is also important from an environmental perspective (Verbič et al., 2019), but so is genetic improvement. In general, about half of phenotypic improvement is due to management and half due to selection (Dekkers & Hospital, 2002).

The ownership of the data that drives dairy improvement is a matter of discussion in many breeding programs. When data collection is subsidized, the data are usually free for management and genetic improvement. Finding optimal frequency of recording for management and genetic improvement in such systems is crucial for optimal return on investment. When data collection is paid by producers, they can be available to breeding organizations for free or at a cost. Depending on the cost breeding organizations can save some resources by purchasing smaller number of repeated phenotypes per individual and genotype more selection candidates instead.

While genotype data is currently largely used for selection, the same data could also support management (predicting feed requirements, disease liability, etc.). Therefore, evaluating the value of phenotype and genotype data is complex and beyond the scope of this study. One possible way forward would be to compare variance between herd-test day effects and genetic variance to contrast the value of managing production and genetic gain in addition to comparing phenotypic and genetic trends (Dekkers & Hospital, 2002).

In practice, test day records are also used to compute the 305-day milk yield (International Committee for Animal Recording, 2020). The longest sampling interval tested in our study and still approved by ICAR was nine weeks, which yielded five records per lactation. Previous studies showed that the correlation of predicting 305-day milk using five records per lactation with using either weekly or eleven records per lactation was between 0.98 and 0.99 (Pool and Meuwissen, 1999; Berry et al. 2005). However, some studies showed that prediction using less than eleven records can yield substantial bias (Gantner et al., 2008)

#### 4.5.2 Single additive trait

We simulated milk yield as a single polygenic trait with additive genetic as well as herd, permanent environment, and residual environmental effects. We did not simulate nor account for non-additive genetic effects that also affect dairy performance (Fuerst & Sölkner, 1994; Ertl et al., 2014; Jiang et al., 2017). We note that we simulated permanent environment effects, which capture non-additive genetic effects and other individual specific environmental effects. We also simulated milk yield in different lactations as a single trait with constant heritability, whereas genetic correlation between different lactations and through the lactation is not unity (Meyer, 1984; Swalve & Vleck, 1987; Dong & van Vleck, 1989). If genetic correlation is less than one, the repeatability of the phenotype decreases and the value of a repeated record and its contribution to accuracy diminishes.

We simulated a trait with heritability of 0.25, since the majority of production traits recorded repeatedly show intermediate heritability. Previous studies also provide insights in how changing the heritability of the phenotype would affect the results. On one hand, at a lower heritability we would need more females in the training population until the contribution of additional female was negligible (Gonzalez-Recio et al., 2014). On the other hand, genomic selection is less affected by the heritability than conventional selection and hence more beneficial for traits with low heritability (Lillehammer et al., 2011; García-Ruiz et al., 2016).

#### 4.5.3 Genomic selection of females

We did not use genomic selection for females, nor did we use reproductive technologies such as sexing semen or embryo transfer. This would further decrease the generation interval, increase selection intensity on female side, and in turn increase genetic gain of genomic scenarios even more (Pryce et al., 2010; García-Ruiz et al., 2016). Such an implementation of genomic selection requires only a minor modification of the design used in this study - genotyping heifers instead of first-parity cows. However, reproductive technologies require a larger modification and investment. Some of the scenarios saved resource and could invest into these technologies.

## 5 Author Contributions

JO designed the study, ran the simulation, analyzed the data, interpreted the results, and wrote the manuscript. JJ participated in study design, results interpretation, and manuscript revision. GG has initiated and supervised the work and contributed to all its stages.

## 7 Acknowledgments

The authors acknowledge I. Pocrnić (The Roslin Institute, University of Edinburgh) for his help in interpreting the results and comments on the manuscript.

## Supplementary Material

### 1. Supplementary Figures and Tables

#### a. Supplementary Tables

**Table S1.** Accuracy of conventional and genomic selection with varying number of phenotypes and phenotyped animals. NoRec = Number of phenotypic records per lactation, NoDaughters = number or daughters per sire, r_sire_ = accuracy for sires, r_rows_ = accuracy for cows, r_non-pheno_ = accuracy for non-phenotyped animals, NoPhenoCows = number of phenotyped cows, NoPhenoTotal = total number of phenotypes (number of phenotypes per lactation times the number of phenotyped cows).

| NoPheno | NoDaughters | $r_{\text{sires}}$ | $r_{\text{cows}}$ | $r_{\text{non-phenotype}}$ | NoPhenoCows | NoPhenoTotal |
| --- | --- | --- | --- | --- | --- | --- |
| **Conventional selection, 100 sires** |  |  |  |  |  |  |
| Variable resources for phenotyping |  |  |  |  |  |  |
| 1 | 100 | 0.93 | 0.62 | 0.56 | 10,000 | 10,000 |
| 2 | 100 | 0.96 | 0.70 | 0.59 | 10,000 | 20,000 |
| 5 | 100 | 0.97 | 0.81 | 0.64 | 10,000 | 50,000 |
| 10 | 100 | 0.98 | 0.89 | 0.66 | 10,000 | 100,000 |
| Fixed resources for phenotyping |  |  |  |  |  |  |
| 1 | 1000 | 0.99 | 0.63 | 0.59 | 100,000 | 100,000 |
| 2 | 500 | 0.99 | 0.71 | 0.61 | 50,000 | 100,000 |
| 5 | 200 | 0.99 | 0.82 | 0.64 | 20,000 | 100,000 |
| 10 | 100 | 0.98 | 0.89 | 0.66 | 10,000 | 100,000 |
| **Genomic selection** |  |  |  |  |  |  |
| Variable resources for phenotyping |  |  |  |  |  |  |
| 1 | - | - | 0.62 | 0.56 | 10,000 | 10,000 |
| 2 | - | - | 0.70 | 0.63 | 10,000 | 20,000 |
| 5 | - | - | 0.81 | 0.71 | 10,000 | 50,000 |
| 10 | - | - | 0.89 | 0.76 | 10,000 | 100,000 |
| Fixed resources for phenotyping |  |  |  |  |  |  |
| 1 | - | - | 0.63 | 0.93 | 100,000 | 100,000 |
| 2 | - | - | 0.71 | 0.90 | 50,000 | 100,000 |
| 5 | - | - | 0.82 | 0.84 | 20,000 | 100,000 |
| 10 | - | - | 0.89 | 0.76 | 10,000 | 100,000 |

**Table S2.**
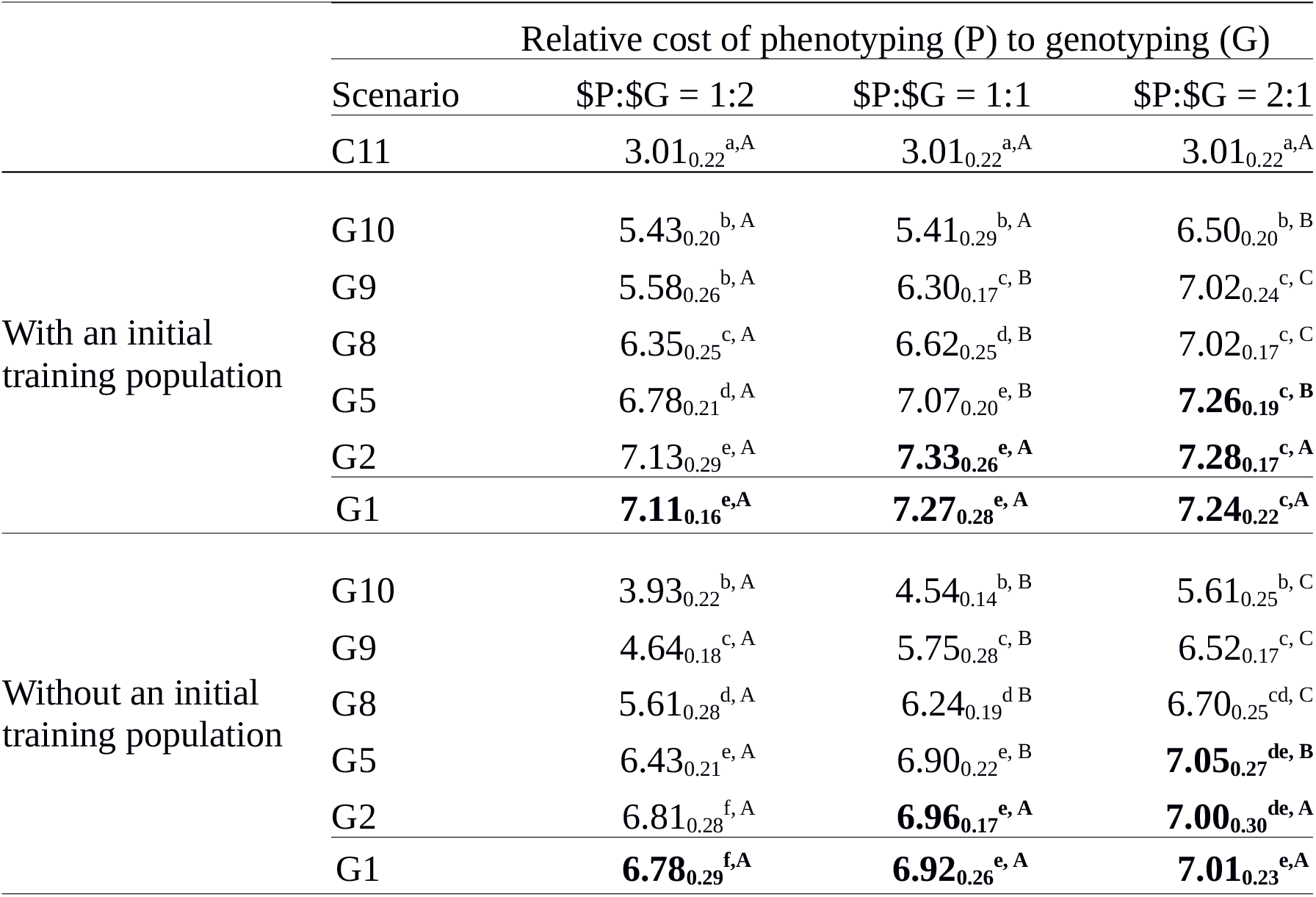
Genetic gain by scenario, relative cost of phenotyping to genotyping ($P:$G), and availability of an initial training population. The table presents the means and standard deviations (subscript) across 10 replicates for the conventional (C) and genomic (G) scenarios, with numbers indicating the number of phenotype records per lactation. For the $P:$G we compared the cost of 11 phenotype records per lactation to the cost of one genotype. The scenarios in bold did not spend all the available resources. Lower-case letters denote statistically significant differences between scenarios within the same $P:$G and upper-case letters between different $P:$G within the same scenario.

**Table S3.** Intensity of sire selection by scenario and relative cost of phenotyping to genotyping ($P:$G). PA = parent average, gEBV = genomic breeding value. We are presenting the bivariate intensities for a two-stage selection. The pre-selection step selects the animals with the highest parent average out of all available new born males to send into testing (progeny of genomical). The selection step selects the sires with the highest breeding values out of all tested males to use in artificial insemination. The scenarios are named C/G for conventional/genomic with numbers indicating the number of phenotype records per lactation. For the $P:$G we compared the cost of 11 phenotype records per lactation to the cost of one genotype.

|  |  | Relative cost of phenotyping (P) to genotyping (G) |  |  |  |
| --- | --- | --- | --- | --- | --- |
| | | Scenario | \$P:\$G = 1:2 | \$P:\$G = 1:1 | \$P:\$G = 2:1 |
|  |  | C11 | 0.10 | 0.10 | 0.10 |
| With an initial<br>training population |  | G10 | 0.17 | 0.27 | 0.37 |
|  |  | G9 | 0.28 | 0.39 | 0.55 |
|  |  | G8 | 0.38 | 0.49 | 0.68 |
|  |  | G5 | 0.59 | 0.77 | 0.91 |
|  |  | G2 | 0.77 | 0.91 | 0.98 |
|  |  | G1 | 0.83 | 0.92 | 0.99 |
| Without an initial<br>training population |  | G10 | 0.17 | 0.25 | 0.36 |
|  |  | G9 | 0.28 | 0.43 | 0.56 |
|  |  | G8 | 0.37 | 0.55 | 0.71 |
|  |  | G5 | 0.58 | 0.77 | 0.91 |
|  |  | G2 | 0.76 | 0.89 | 0.98 |
|  |  | G1 | 0.83 | 0.94 | 0.99 |

**Table S4.**
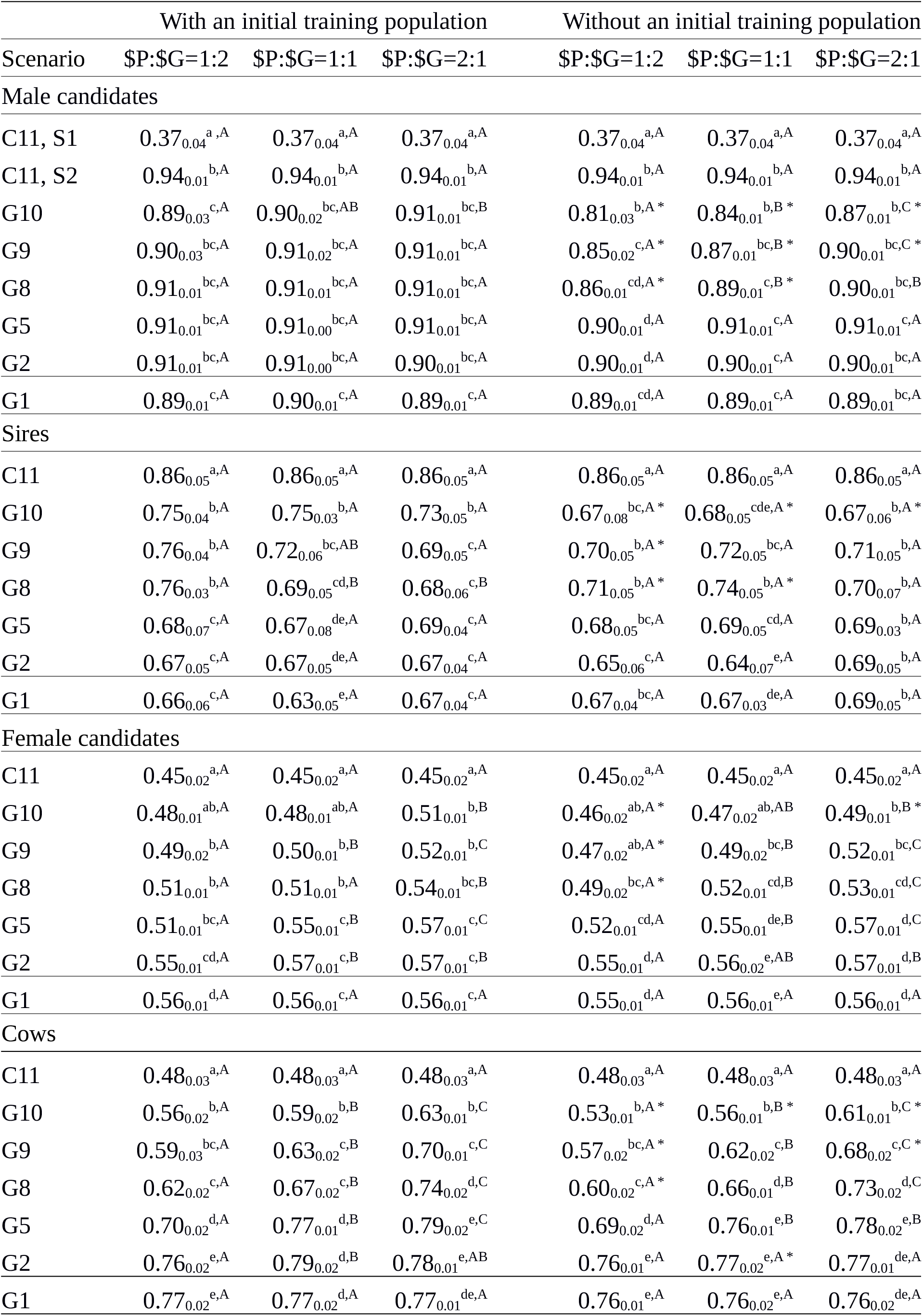
Selection accuracy by scenario, relative cost of phenotyping to genotyping ($P:$G), and the availability of an initial training population. The table presents the means and standard deviations (subscript) across 10 replicates for the conventional (C) and genomic (G) scenarios, with numbers indicating the number of phenotype records per lactation. For the $P:$G we compared the cost of 11 phenotype records per lactation to the cost of one genotype. Conventional selection implemented two-stage selection for males, hence we present the accuracy of pre-selection for progeny testing (S1) and the accuracy sire selection (S2). In genomic scenarios the male candidates were genotyped and non-phenotyped. We also present the accuracy for sires currently used in artificial insemination (sires), for non-genotyped and non-phenotyped females (female candidates), and for all active phenotyped cows and bull dams (cows). Lower-case letters denote statistically significant differences between scenarios within the same $P:$G and upper-case letters between different $P:$G within the same scenario. Stars denote statistically significant difference between corresponding scenarios with and without an initial training population.

#### b. Supplementary Figures

**Figure S1.**
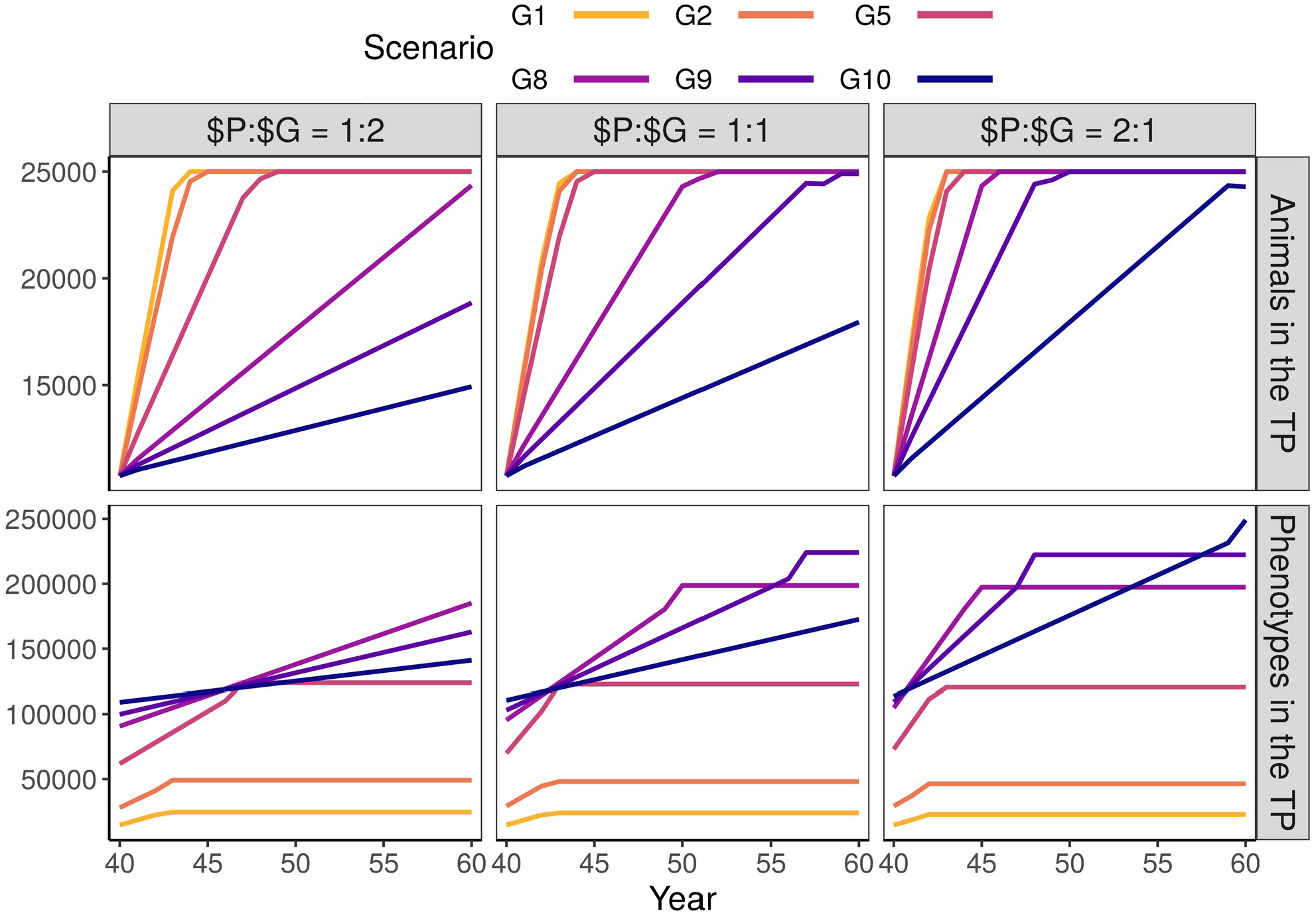
The number of animals and repeated phenotypes in the training population by scenario and relative cost of phenotyping to genotyping ($P:$G) with an initial training population (TP). The scenarios are named C/G for conventional/genomic with numbers indicating the number of phenotype records per lactation. For the $P:$G we compared the cost of 11 phenotype records per lactation to the cost of one genotype. In our simulation, scenarios traded repeated phenotype records for genotypes. Hence, the scenarios with the largest training population collected the least repeated records. These were also the scenarios that achieved the highest genetic gain.

